# Complex functional phenotypes of NMDA receptor disease variants

**DOI:** 10.1101/2022.07.01.498520

**Authors:** Gary J Iacobucci, Beiying Liu, Han Wen, Brittany Sincox, Wenjun Zheng, Gabriela K. Popescu

**Author notes:** equal contributions (Gary J Iacobucci, Beiying Liu). Senior coauthors (Wenjun Zheng, Gabriela K. Popescu). To whom correspondence should be addressed Gary J Iacobucci: Department of Biochemistry, Jacobs School of Medicine and Biomedical Sciences, University at Buffalo, Buffalo, NY 14203 Gabriela K Popescu: Department of Biochemistry, Jacobs School of Medicine and Biomedical Sciences, University at Buffalo, Buffalo, NY 14203; Wenjun Zheng: Department of Physics, College of Arts and Sciences, University at Buffalo, SUNY, Buffalo NY 14260.

## Abstract

NMDA receptors have essential roles in the physiology of central excitatory synapses and their dysfunction causes severe neuropsychiatric symptoms. Recently, a series of genetic variants have been identified in patients, however, functional information about these variants is sparse and their role in pathogenesis insufficiently known. Here we investigate the mechanism by which two GluN2A variants may be pathogenic. We use molecular dynamics simulation and single-molecule electrophysiology to examine the contribution of GluN2A subunit-residues, P552 and F652, and their pathogenic substitutions, P552R and F652V, affect receptor functions. We found that P552 and F652 interact during the receptors’ normal activity cycle; the interaction stabilizes receptors in open conformations and is required for a normal electrical response. Engineering shorter side-chains at these positions (P552A and/or F652V) caused a loss of interaction energy and produced receptors with severe gating, conductance, and permeability deficits. In contrast, the P552R sidechain resulted in stronger interaction and produced a distinct, yet still drastically abnormal electrical response. These results identify the dynamic contact between P552 and F652 as a critical step in the NMDA receptor activation, and show that both increased and reduced communication through this interaction cause dysfunction. Results show that subtle differences in NMDA receptor primary structure can generate complex phenotypic alterations whose binary classification is too simplistic to serve as a therapeutic guide.

**Main findings:**

- Two NMDA receptor residues whose substitution results in encephalopathies, were found to form new interactions during activation, and the energy provided by this interaction is required for normal receptor gating.
- Experimental substitutions of these residues that change the strength of their interaction reduce the receptor open probability, unitary conductance, and calcium permeability.
- Receptors with variations at these positions identified in patients display a broad range of both gain- and loss-of-function changes depending on the stimulation protocol.

## Introduction

N-methyl-d-aspartate (NMDA) receptors are glutamate-gated excitatory channels with critical roles in the normal development and function of the nervous system. They are principal mediators of synaptic formation, maturation, and plasticity throughout the life span. In turn, both their insufficient and excessive activation have been long known to cause severe neuropathologies. More recently, gene sequencing approaches in patients with neuropsychiatric disorders have identified alterations in the primary structure of *GRIN* genes, which encode NMDA receptor subunits ^1–7^. To help with patient stratification and therapy development, several publicly-available databases centralize information on the rapidly increasing number of clinically reported variants ^8^. This aggregation has made apparent several challenges that, at present, obscure the disease mechanism of these variants.

First, rather than being specific and localized to specific genetic/structural regions, the identified genetic alterations are diverse and widely spread over the entire length of all seven NMDA receptor subunits. Second, a direct correlation between the primary structure of NMDA receptors subunits and their functional output remains elusive. Lastly, how NMDA receptor responses affect the normal physiology of the central nervous system, and specifically which of their properties are important at a particular time and place, is only superficially understood. To date more than 4,000 *GRIN* variants have been identified in human populations. About half of these have no reported clinical phenotype and are currently classified as “benign.” The remainder display clinical features ranging in impact from mild to severe and include developmental delay, epilepsy, schizophrenia, intellectual disability, autism spectrum disorders, attentionLJdeficit and hyper-activity disorders, visual impairment, hypotonia, speech disorders, movement disorders, and microcephaly ^9–11^. In part, this pleiotropy likely reflects the receptor’s diverse and dynamic contributions to ongoing normal functionality of synapses, neurons, and circuits. It also reflects the complex and insufficiently understood relationship between the receptor’s primary structure and its healthy operation.

NMDA receptors are obligate heterotetramers that assemble from two obligatory GluN1 subunits, encoded by *GRIN1*, and a collection of two GluN2 or GluN3 subunits, encoded by *GRIN2A-D* and *GRIN3A-B*, respectively. Consistent with its required role for NMDA receptor assembly and expression, GluN1 subunits are widely expressed across brain regions and developmental stages; and animals lacking the GluN1 perish at birth due to respiratory failure ^12, 13^. In contrast, the expression of GluN2 and GluN3 subunits is developmentally and regionally controlled, and animals lacking these subunits have severe but non-lethal phenotypes ^14^. Disease-associated variants have been identified in all eight NMDA-receptor encoding genes, attesting for the critical and non-redundant roles of individual subunits.

Tetrameric NMDA receptors are large (∼4,500 residues) transmembrane proteins. About two-thirds of residues are extracellular and are organized into two layers, each consisting of four globular domains. The membrane-distal N-terminal layer forms modulator-binding sites and influences the channel open probability, but is dispensable to agonist-dependent activation ^15, 16^. Likely, mutations in this layer affect the receptor’s sensitivity to allosteric modulators. The membrane proximal ligand-binding layer consists of four globular domains, which form binding sites for the physiologic co-agonists glutamate and glycine. Agonists stabilize a more compact set of receptor conformations and energetically couple with increased mechanical tension in the three short linkers that connect each ligand-binding module to one of three transmembrane helices (M1, M3, and M4). Together with a short M2 helix, which inserts in the membrane as a P-loop from the cytoplasmic surface, transmembrane helices surround the cation-permeable pore, and residues on the M3 helix opposite to the cytoplasmic surface form the agonist-controlled gate. The ligand-binding and transmembrane domains, together with the linkers that connect them form the core of the NMDA receptor channels in that they are required and sufficient for their defining function as glutamate-gated excitatory channels. The cytoplasmic domain is large; it represents about a third of receptor mass; and although it is dispensable for glutamate-gated currents ^17^, animals lacking this domain are not viable ^18^. This observation, together with the lack of a specific associations between disease-related variants and the receptor’s various structural domains, indicate critical roles for all receptor domains in the normal physiology of the central nervous system.

Although identified simply as glutamate-gated excitatory channels, NMDA receptors are complex multifunctional proteins. In addition to binding glutamate, their typical activation cycle also includes interactions with a host of organic and inorganic ions and molecules as diverse as inorganic cations such as protons, magnesium, and calcium, and a host of organic molecules, which include glycine, polyamines, steroids, and several proteins. In turn, the residues that form these external ligand-binding sites are internally connected to the channel gate through complex networks of allosteric interactions. In effect, these internal interaction networks transform the binding energy contributed by ligands into conformational changes that alter the receptor’s overall function. Therefore, structural variations as minor as a single-residue substitution can affect the intensity and duration of its glutamate-elicited ionic current by changing how ligands bind or how the binding energy is transmitted to the gate. Of the patient-derived variants that have been classified as pathogenic, fewer than half have been examined functionally, and even fewer have a proposed mechanism ^1, 5, 6, 19–29^. In part, this is because the activation mechanism of NMDA receptors is complex and insufficiently understood ^30, 31^ making correlation between *in vitro* characterization and *in vivo* behavior difficult ^32, 33^. To begin to explain how NMDA receptor mutations alter receptor functions and how to restore pharmacologically their normal operation, it is necessary to outline the mechanism by which individual residues contribute to the receptor’s normal operation and how their substitution alters receptor responses.

In a previous study, we used molecular dynamics simulations and identified pairs of interacting residues whose strength of interaction changes during activation. Among these, several corresponded to sites where variations are pathogenic ^34^. Specifically, GluN2A F652, which has been associated with epilepsy, contacted in a state-dependent manner GluN1 R801, for which no variation has yet been reported, and GluN2A P552, which is also associated with epilepsy. We hypothesized that mutations at either GluN2A F652 or GluN2A P552 may work through the same mechanism, namely by changing their interaction which is intrinsic to the activation sequence. Here, we report evidence for the direct chemical coupling between F652 and P552 during receptor opening and show that mutations at these sites, regardless of whether they weaken or strengthen the native interaction, result in severe deficits in receptor open probability. Notably, these deficits produced distinct functional phenotypes, which could not be predicted by simple functional characterization.

## Methods

### Cell culture and molecular biology

HEK293 cells (ATCC CRL-1573) at the passages 25 – 32 were maintained in Dulbecco’s Modified Eagle Medium with 10% Fetal Bovine Serum and 1% glutamine. Cells were incubated at 37°C in 5% CO_2_ and 95% atmospheric air. Prior to experiments, cells were transfected with rat GluN1-1a (U08262.1), GluN2A (M91561.1), or mutants as indicated, and GFP cDNA at a 1:1:1 ratio using polyethylenimine ^35^. The transfected cells were grown for 24 – 48 h in medium supplemented with 10 mM Mg^2+^ to prevent excitotoxicity. All mutations were introduced by using the QuickChange method (Stratagene, La Jolla, CA) and verified by DNA sequencing.

### Electrophysiology

Stationary single-channel currents were recorded with the cell-attached patch-clamp technique at holding potential +100 mV. Borosilicate pipettes (15 – 25 MΩ) contained (extracellular, in mM): 150 NaCl, 2.5 KCl, 0.1 EDTA, 10 HEPBS, 0.1 glycine, 1 glutamate, pH 8.0 (adjusted with NaOH). Currents were amplified and filtered at 10 kHz (Axopatch 200B; 4-pole Bessel), sampled at 40 kHz (PCI-6229, M Series card, National Instruments, Austin, TX) and written into digital files with QuB acquisition software (University at Buffalo, Buffalo, NY). For the experiments with Ca^2+^, patches were held at potentials between −100 to −20 mV, in 20 mV increments.

Macroscopic currents were recorded with the whole-cell patch-clamp technique using borosilicate pipettes (2 – 4 MΩ) containing (intracellular, in mM): 135 CsCl, 33 CsOH, 0.5 CaCl_2_, 2 MgCl_2_, 11 EGTA, 10 HEPES, pH 7.4 (adjusted with CsOH) and clamped at −70 mV. Extracellular solutions contained (in mM): 150 NaCl, 2.5 KCl, 0.5 CaCl_2_, 0.01 EDTA, 0.1 glycine, 1 glutamate, 10 HEPBS, pH 8.0 (NaOH). Currents were amplified and filtered at 2 kHz (Axopatch 200B; 4-pole Bessel), sampled at 5 kHz (Digidata, 1440A) and written into digital files with pClamp 10 acquisition software (Molecular Devices, Sunnyvale, CA). Clamped cells were extracellularly perfused with solutions using BPS-8SP solenoid-valve perfusion system (ALA Scientific Instruments, Westbury, NY). Free Ca^2+^ concentrations were calculated with the software MAXC (www.maxchelator.stanford.edu).

To evaluate receptor permeability to Ca^2+^, we determined reversal potential (E_rev_) of macroscopic currents by applying voltage ramps from −100 mV to +60 mV over 4 sec on glutamate-elicited steady state currents in several external Ca^2+^ concentrations. Currents were leak-subtracted using currents elicited with voltage ramp in the absence of glutamate. Liquid junction potentials were measured in each condition using the K^+^ salt-bridge method ^36^. E_rev_ was calculated by a linear fit between −20 to +20 mV using the current-voltage data corrected for leak current and liquid junction potentials The magnitude of the Ca^2+^-induced shift in E_rev_ relative to Ca^2+^-free conditions was related to Ca^2+^ permeability using the Lewis equation ^37^:

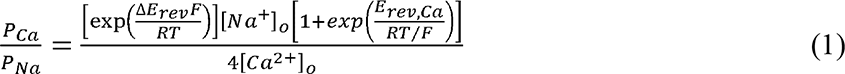

### Data Analysis

Single-channel traces were corrected for noise artifacts and baseline drift and idealized in QuB software with the segmental-k-means (SKM) algorithm after applying a 12 kHz digital low-pass filter ^38^. All subsequent analyses of idealized records were done in QuB with the maximum interval log-likelihood (MIL) algorithm after imposing a conservative dead time (75 μs) ^39^. Rate constants were estimated by fitting a model with five closed and two open states (5C2O) directly to the idealized data ^40^. To determine the duration-weighted rate constants to be used for macroscopic current simulation, the model was fit globally to data pooled across all patches. Bursts of activity were defined as openings separated by closures shorter than a critical duration (τ_crit_) calculated to exclude desensitized periods. Once defined, bursts were extracted and analyzed separately.

Macroscopic currents were analyzed in Clampfit 10.7 (Molecular Devices). Steady-state (I_ss_) current amplitude was measured as the average current amplitude at the end of a 5 or 10 sec pulse of glutamate at the indicated concentration. For dose-response analysis, I_ss_ measured at each dose was normalized to the max I_ss_ measured in 0.1 mM glutamate. To estimate the effective glutamate binding rate for receptors containing GluN2A^P552R^, time constants were determined by fitting a single-component exponential function to estimate the effective activation time-course (τ_rise_). The resulting time constants were plotted against the inverse of agonist concentration as conventional for a two-site ligand binding model. K_on_ was determined from the slope of a linear fit to this data.

### Simulations

Macroscopic current traces were calculated as the time-dependent accumulation of receptors in open states using kinetic models and unitary amplitudes derived from single-molecule recordings. All receptors occupied initially a glycine-bound glutamate-free resting state connected to fully liganded state C3 with the rate constants determined previously for wild-type receptors ^41^.

Simulations were performed in MATLAB 2017a (Mathworks) using the built-in matrix exponential function, *expm*. We used the rate constants derived in QuB to construct a matrix (*A*) of *n* x *n* size where *n* is the number of states in the model and each element is the rate constant value between the corresponding states. A deterministic simulation of the occupancy of all states with time resolution, *dt*, was performed by solving iteratively:

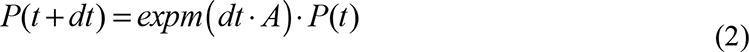

The final macroscopic current amplitude was calculated by summing the occupancies of both open states in the model at each time point. Total charge transfer was calculated as the integral of the resulting current waveform.

### Molecular Dynamics Simulations

The core GluN1/GluN2A structural model used in this study was generated by homology modeling followed by targeted-molecular dynamics simulations ^34, 42^. Briefly, the GluN1/GluN2A homology model was generated with SWISS-MODEL using the GluN1/GluN2B crystal structure (4tlm) ^43^ to leverage its superior resolution of linker residues and the several reported distinct conformations which can be used as templates for targeted molecular dynamics (MD) simulations ^44^. Targeted MD simulations were performed with NAMD V2.9b using a putative active-state structure as the target conformation ^44^. In all MD trajectories, only the last 150 ns period was used for energy analysis and the last 10 ns period was used for HOLE calculation ^45^. We estimated inter-residue Van der Waals energies with the NAMDEnergy module in the VMD program ^46, 47^.

### Statistics

All results are presented as means with the associated standard errors (mean ± sem). Statistical significance of differences was evaluated with the paired or unpaired *t* test, as appropriate. Differences were considered significant for *p* < 0.05.

## Results

### Intrasubunit coupling between residues in the GluN2A D1-M1 linker and M3 helix

In previous work, we used targeted MD simulation of a core GluN1/GluN2A construct lacking both N- and C-terminal domains (N1_ΔNΔC_/N2A_ΔNΔC_) to identify pairs of side-chains predicted to engage in state-specific interactions during NMDA receptor activation ^34^ (Figure 1a). Among these, we prioritized for functional testing residues for which naturally occurring variants were suspected as the cause of behavioral dysfunctions, as annotated in contemporary databases ^48^. We found that residues located on the short segments linking LBD with TMD, rather than acting simply as non-specific mechanical springs ^49^, form specific chemical interactions, which catalyze the channel-opening reaction, and are critical for the receptor’s physiological function. Motivated by this new insight, we considered additional residue pairs flagged by the MD simulation as potentially forming state-dependent chemical contacts.

**Figure 1.**
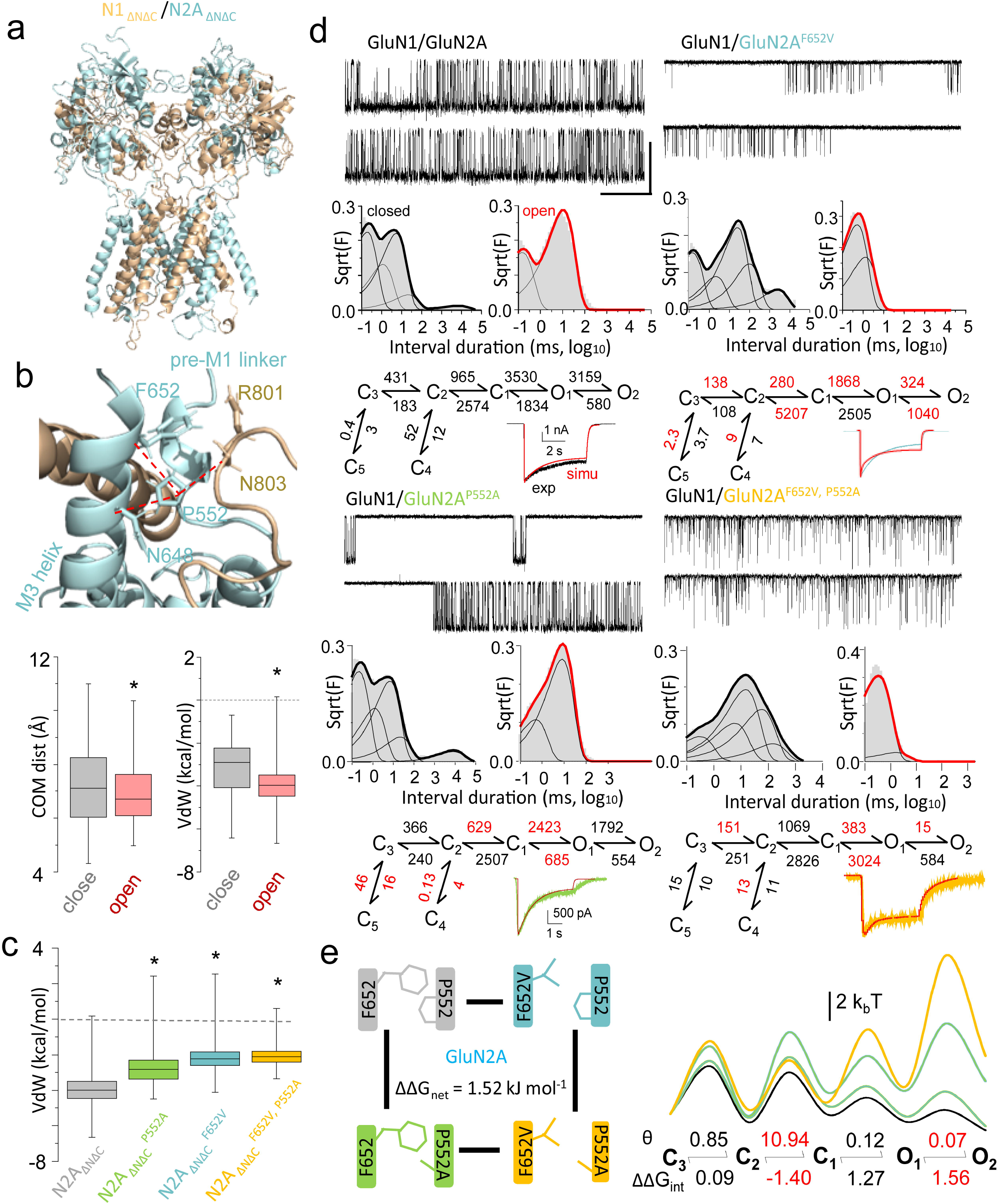
Two residues important for normal neurological function interact during NMDA receptor gating. (**a**) Structural model of a core GluN1/GluN2A receptor lacking NTD and CTD (N1_ΔNΔC_, tan; N2A_ΔNΔC_, cyan). (**b**) Within the GluN2A subunit, the interaction between P552 and F652 *(top*) is activity-dependent as indicated by smaller center of mass distance (COM) (*bottom left*), and stronger Van der Waals contact energy (VdW) (*bottom right*), in open versus closed conformations. \**p* < 0.05 Kolmogorov-Smirnoff test. (**c**) Substitutions at both P552 and F652 change the VdW contact energy between these residues in structural models of open receptors, consistent with the disruption of a gating-favorable interaction. \**p* < 0.05 Kolmogorov-Smirnoff test. (**d**) Side chains of both P552 and F652 contribute to the gating kinetics of full-length NMDA receptors; substitutions at these sites (P/A and F/V) changed the pattern of current recorded from individual receptors, the distribution of closed (black) and open (red) intervals, and the gating rate constants calculated with the indicted state models. Macroscopic current responses to pulses (5 s) of glutamate (1 mM) predicted by each model (red) are shown superimposed with experimentally recorded whole-cell currents (green and yellow). \**p* < 0.05 two-tailed t-test. (**e**, *left*) Diagram of the thermodynamic cycle used to calculate the coupling energy between P552 and F652 using the rates illustrated in panel d. (*right*) Energy landscapes calculated for the gating reactions of individual receptors illustrate increased barriers to activation in receptors with mutations at disease-associated residues.

We noted that P552, which resides on the GluN2A D1-M1 linker, contacted four proximal residues on the same subunit: F652 and N648 on the M3 helix, and R801 and N803 on the D2-M4 linker (Figure 1b). Of these putative interactions, we chose to examine in more depth the relationship between GluN2A-P552 and GluN2A-F652 for two reasons. First, along the closed-to-open trajectory of the MD simulation, their side chains moved closer together as measured by center of mass (COM) distance, and formed more favorable Van der Waals (VdW) interactions in the open state relative to the initial conformation (Figure 1b, Figure S1). Second, variants with substitutions at these positions have been identified in patients with epilepsy and intellectual disability, suggesting that they have a critical role in receptor’s biological function in the central nervous system ^4, 5^. As a preliminary step in our study, we used the open structural model we generated previously ^34^ to ask whether removing the side-chains of GluN2A-P552 and GluN2A-F652 would affect the VdW contact energy between these residues. Results for receptors containing GluN2A-P552A or GluN2A-F652A showed significant change in VdW contact energy consistent with the disruption of a gating-favorable interaction (Figure 1c, Figure S1). We hypothesized that state-dependent interactions between these residues represents a critical step in the opening sequence, which when disrupted cause pathological electrical signaling.

We proceeded to test this hypothesis by measuring the strength of the interaction between P552 and F652 in full-length receptors expressed in HEK 293 cells using double-mutant cycle analyses ^50^. In this approach, the residues suspected of functional coupling are mutated both individually and together, and the free-energy landscape of the gating reaction is measured for each variant, to estimate the individual and combined effects of the two residues. If the change observed for the double mutant is simply the arithmetic sum of the changes observed for individual mutations, the residues likely make independent contributions to gating; whereas departures from simple addition indicate that the residues interact and the interaction energy contributes specifically to gating. Importantly, when the energy landscape is computed from measurements obtained from single-molecule observations, results inform not only globally about the roles played by the probed residues in the overall activation sequence, but they quantify explicitly their contributions to each gating step.

We recorded equilibrium activity from receptors engineered to contain GluNA^P552A^, GluN2A^F652V^, or GluN2A^P552A, F652V^, in combination with wild-type GluN1-1a as described previously ^51^ (Figure 1d). We could not observe macroscopic of microscopic currents from receptors containing the GluN2A^F652A^ variant (data not shown), and the activity of receptors containing GluN2A^F652V^ was dramatically impaired, suggesting that this position is critical for gating. Records obtained from wild-type receptors and the remaining three variants were processed and used for kinetic modeling to estimate rate constants for each receptor’s activation sequence with the usual methods ^52, 53^. Next, we validated the reaction schemes obtained by comparing the waveform of their predicted macroscopic response with experimentally recorded whole-cell currents (Figure 1d). Based on the satisfactory match between responses predicted with models derived from single-channel data and those recorded directly from cells expressing each variant, we used the models to calculate free-energy landscapes for each receptor, and to estimate the coupling free energy for the double-mutant thermodynamic cycle ^34, 54^ (Figure 1e). Results show a net surplus of 1.52 kJ/mol free energy (ΔΔG_int_) for the double mutant over the entire gating reaction, which indicates that P552 and F652 interact substantially during gating, and the energy generated by this interaction makes an important contribution to the physiologic gating kinetics of NMDA receptors.

Considering in more detail the steps within the activation sequence of each variant, we noted that the interaction between P552 and F652 had the largest impact in the later steps of the gating sequence, and facilitated specifically the C_2_-C_1_ (−1.4 kcal/mol) and O_1_-O_2_ (1.56 kcal/mol) transitions. This result suggests that the contacts between P552 and F652, which are favorable to channel opening, occur after the receptors transitions through pre-open conformations and they serve to stabilize open-gate conformations. This interpretation is consistent with the results from the targeted MD simulation, which predicted a reduction in side-chain distance between these residues during activation, and an increased energetic interaction; whereas side-chain truncation of either or both residues reduced the VdW energy (Figure 1b, c). Together with the results from our thermodynamic analysis these observations support the view that in wild-type receptors, the side-chains of P552 and F652 contribute essentially to the normal opening of NMDA receptors; therefore, preceding glutamate-induced movements in the LBD that bring these residues within interaction distance will catalyze the opening reaction by forming a chemical link that stabilizes the open-pore conformation. A corollary of this finding is that substitutions at these positions that prevent the harnessing of LBD kinetic energy into chemical energy to stabilize the open state, will present gating deficits, as illustrated by the functional analyses for receptors with truncated side chains described above. However, it remains unanswered whether the variants identified in patients affect NMDA receptor function simply because they lack this interaction.

### Disease-associated variants modify interactions between the D1-M1 linker and the M3 helix

Pathogenic variations in GluN2A at P552 and F652 have been identified in patients; specifically, GluN2A^F652V^ (ClinVar: VCV000088733.1) ^5^ and GluN2A^P552R^ (ClinVar: VCV000039663.3) ^4^ (Figure 2a, *left*) are accompanied by an array of neurological dysfunctions. It is therefore important to ascertain whether these specific mutations affect receptor function with the same mechanism as described above for mutants with side-chain truncations. When we introduced these naturally occurring substitutions in the structural model of activated receptors, we observed that they had distinct effects on the VdW energy of interaction. Relative to wild-type receptors, receptors with GluN2A^F652V^ had less favorable interaction energy between P552 and V652, whereas receptors with GluN2A^P552R^ had substantially more favorable interaction energy between R552 and F652 (Figure 2b, Figure S1). Previous studies have already documented that both these mutations have strong effects on gating ^5, 20^. However, the mechanism by which the described changes occur remains unresolved.

**Figure 2.**
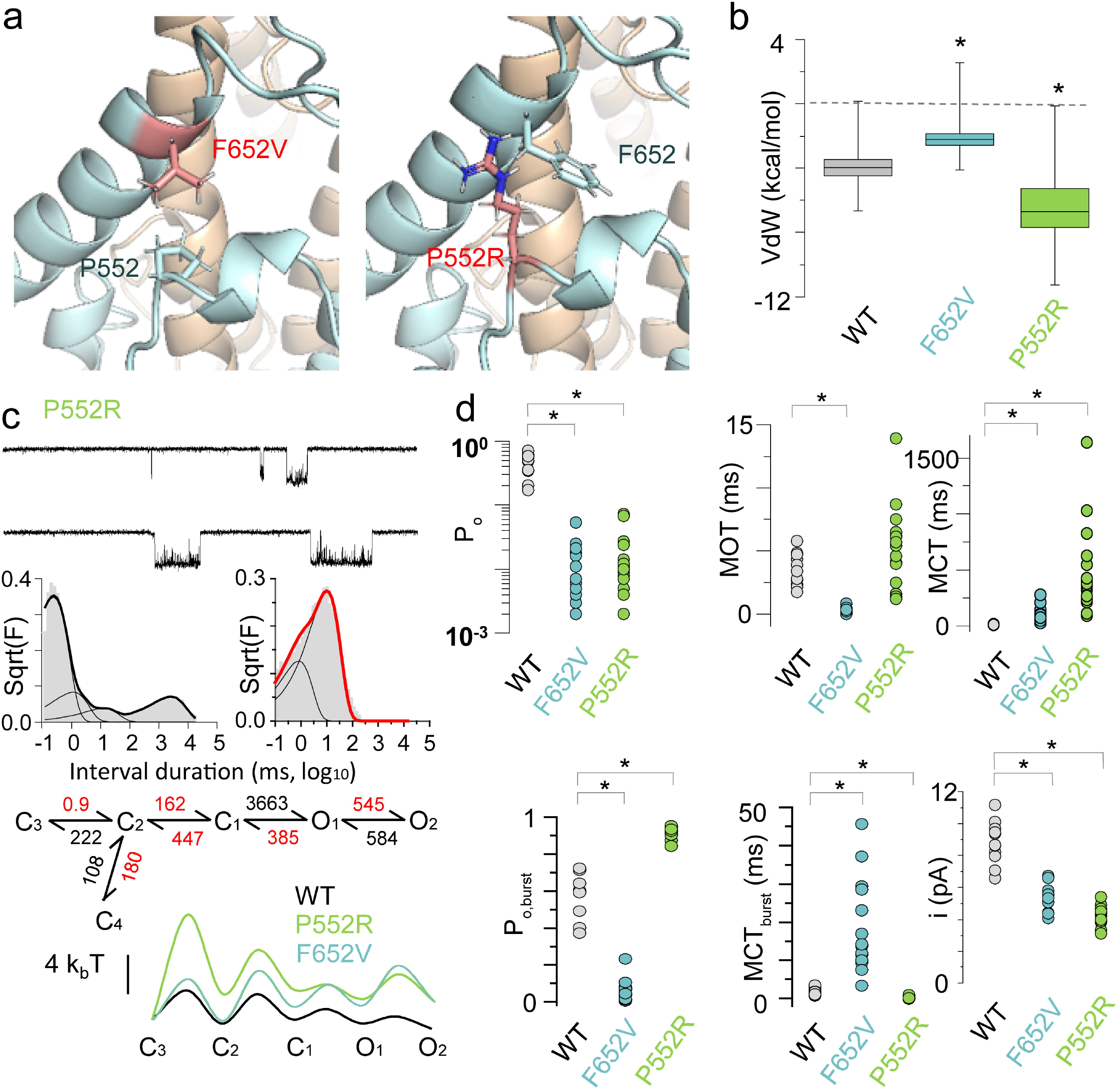
NMDA receptor variants associated with neurological dysfunction display a broad range of gating perturbations. (**a**) Structural models of open NMDA receptors variants illustrate the relative positions of two disease-associated residues. (**b**) Relative to receptors with wild-type residues, the modeled open states of receptors with disease-associated mutations have distinct Van der Waals contact energies (VdW) between residues 552 and 652 of GluN2A (\**p* < 0.05; Kolmo-gorov-Smirnov test). (**c**, *top*) Currents recorded from individual full-length GluN1/GluN2A ^P552R^ receptors (*n* = 16); (*middle*) Dwell-time histograms of closed (left) and open (right) interval durations with superimposed distributions (lines) predicted by the model illustrated below; (*bottom*) Energy landscapes calculated from the kinetic models derived for the indicated receptors. (**d**) Distributions of gating parameters estimated for the indicated full length receptor types: open probabilities (P_o_), mean open (MOT) and mean closed (MCT) durations estimated for entire records or for bursts of activity. (\**p* < 0.05; Student’s *t* test).

To characterize the gating reactions of these two naturally occurring variants we recorded cell-attached currents from patches with a single active receptor and subjected these data to kinetic analyses and modeling (Figure 2c). We observed substantial alterations in the gating profiles of both variants. Notably, for receptors containing GluN2A^P552R^, we could only discern four closed states, rather than the typical five observed for native receptors; and for both variants the transition rates between the kinetic states detected were profoundly altered. Relative to wild-type receptors, the computed free-energy landscapes were substantially elevated for both variants, with kinetic states sitting in shallower wells and being separated by larger barriers. For receptors containing GluN2A^P552R^, the largest transition barrier occurred early in the gating reaction such that, at equilibrium, substantially fewer receptors transitioned into open states; however, the fewer receptors that managed to open, remained open for longer times, being unable to return to states from which agonists could dissociate to terminate the activation reaction (Figure 2c). This mechanism explains well the previously reported changes in macroscopic current, including increased agonist potency and efficacy, slower rise time, and slower deactivation ^5, 20^. However, it is important to note that these macroscopic behaviors can only be observed for the few receptors that happen to open during the observation window (<5 s), whereas the majority of receptors remain electrophysiologically silent. By observing an individual channel over a large period (>30 min) we obtained a more realistic view of the receptor’s energy landscape.

We noted large variability in the kinetic properties of these receptors (Figure 2d; Table S1). Except for their open probability within bursts (P_o,burst_) and their unitary current amplitudes (*i*), both variants had substantially more variable open probability (P_o_), mean open times (MOT), and mean closed times (MCT) relative to wild-type receptors. Specifically, GluN2A^P552R^ had highly variable open durations, such that although longer on average, the difference was not statistically significant relative to wild-type receptors. Notably, the two variants had distinct burst structures, with substantially higher open probability for GluN2A^P552R^ and lower open probability for GluN2A^F652V^.

These functional results are consistent with the predictions from the MD simulation (Figure 2b), where GluN2A^P552R^ displayed VdW interactions more favorable to opening relative to the wild-type residues. This may be explained by the larger sidechain surface area available for contacts. In addition, we observed an electrostatic cation-π interaction between the arginine side chain and the aromatic ring of phenylalanine, which likely contributes further energy to strengthen the interaction between these residues in open receptors. This interpretation is consistent with the observed longer openings and shorter closures durations in bursts for GluN2A^P552R^. In contrast, GluN2A^F652V^ which was predicted to have fewer VdW contacts and less favorable interaction energy with P552 produced shorter openings and longer closures within bursts. We conclude that stronger interactions between residues located on the D1-M1 linker and the M3 helix contribute directly to the stability of open-gate conformations and increase channel open times and open probability within bursts.

### Interactions between residues in the D1-M1 linker and M3 helix modulate receptor permeation

Both GluN2A^F652V^ and GluN2A^P552R^ had lower unitary current amplitudes (Table S1; Figure 2c). Given that our single channel measurements were done with sodium as the main permeant ion to facilitate detection of gating steps ^55^, these measurements do not offer information about possible changes in the Ca^2+^ content of the reduced currents. To determine whether these mutations affected Ca^2+^ permeability, which is a critical aspect of NMDA receptor signals, by measuring the relative permeability of Ca^2+^ to monovalent ions (P_Ca_/P_Na_) using the magnitude of the shift in measured reversal potential (E_rev_) of macroscopic currents (Figure 3a, b). We elicited whole-cell currents from each mutant and applied a voltage ramp protocol on the steady-state current to measure the reversal potential (E_rev_) at several external Ca^2+^ concentrations.

**Figure 3.**
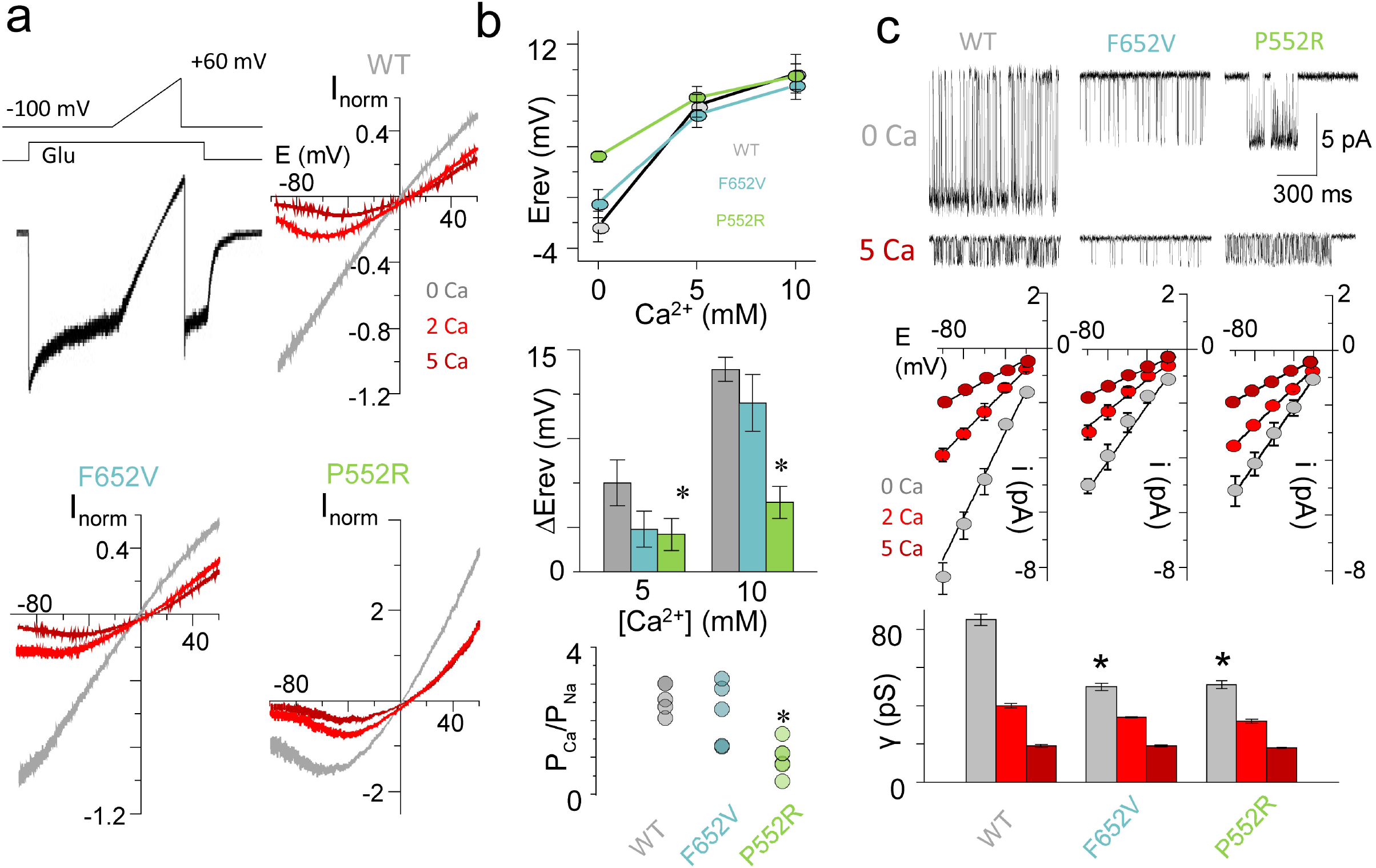
NMDA receptor variants associated with neurological dysfunction display changes in conductance and permeability. (**a**) *Top*, Whole-cell current trace recorded in response to Glu application (1 mM) illustrates change in steady-state current amplitude during a ramp in the membrane potential. *Bottom*, Macroscopic current-voltage relationships measured from recordings as in panel a. (**b**) Ca^2+^ permeability properties inferred from macroscopic current recordings as in a. (**c**) Top, Unitary current traces recorded from cell-attached patches with +100 mV applied potential and external Ca^2+^ as indicated. *Bottom*, Unitary current-voltage relationships for the indicated receptors and summary of results. (\**p* < 0.05; Student’s *t* test relative to WT).

We found relative to wild-type receptors (P_5Ca_/P_Na_ = 2.7 ± 0.5, P_10Ca_/P_Na_ = 2.7 ± 0.2, n = 5), channels harboring GluN2A^P552R^ exhibit reduced Ca^2+^ permeation in 5 mM (P_5Ca_/P_Na_ = 1.2 ± 0.1, n = 10, p = 0.04) and 10 mM Ca^2+^ (P_10Ca_/P_Na_ = 1.1 ± 0.2, n = 6, p = 1.2E-4). By contrast, channels harboring GluN2A^F652V^ exhibit similar Ca^2+^ permeation in 5 mM (P_5Ca_/P_Na_ = 1.7 ± 0.4, n = 6, p = 0.17) and 10 mM Ca^2+^ (P_10Ca_/P_Na_ = 2.3 ± 0.4, n = 5, p = 0.35) compared to wild-type.

Further, we measured for each receptor the slope unitary conductance in several external Ca^2+^ concentrations and calculated the amount of Ca^2+^-dependent reduction in unitary conductance (Ca^2+^ block) as described previously ^56^ (Figure 3c). Wild-type receptors exhibit high Na^+^ unitary conductance (γ_Na_ = 85.5 ± 3.0, *n* = 5) which decreases with increasing extracellular Ca^2+^ concentrations (γ_2Ca_ = 39.7 ± 1.1, *n* = 7, *p* = 1.8E-4; γ_5Ca_ = 18.9 ± 0.7, *n* = 5, *p* = 1.2E-4). This corresponds with strong Ca^2+^ block (γ_Na_/γ_2Ca_ = 0.46, 95% CI = 0.45 – 0.48; γ_Na_/γ_5Ca_ = 0.22, 95% CI = 0.21 – 0.23). Receptors harboring GluN2A^P552R^ showed decreased Na^+^ conductance (γ_Na_ = 50.5 ± 1.6, *n* = 6, *p* = 2.5E-4 to WT), which decreases with increases extracellular Ca^2+^ concentrations (γ_2Ca_ = 31.6 ± 1.2, *n* = 7, *p* = 4.3E-4 to WT; γ_5Ca_ = 18.5 ± 0.3, *n* = 8, *p* = 0.60 to WT). This corresponds with reduced Ca^2+^ block compared to wild type receptors (γ_Na_/γ_2Ca_ = 0.63, 95% CI = 0.60 – 0.65; γ_Na_/γ_5Ca_ = 0.37, 95% CI = 0.35 – 0.38). Similarly, receptors harboring GluN2A^F652V^ showed decreased Na^+^ conductance (γ_Na_ = 50.4 ± 2.1, *n* = 6, *p* = 9.5E-5 to WT), which decreases with increases extracellular Ca^2+^ concentrations (γ_2Ca_ = 33.7 ± 0.3, *n* = 13, *p* = 1.4E-3 to WT; γ_5Ca_ = 18.7 ± 0.5, *n* = 6, *p* = 0.83 to WT). This corresponds with reduced Ca^2+^ block compared to wild type receptors (γ_Na_/γ_2Ca_ = 0.67, 95% CI = 0.66 – 0.69; γ_Na_/γ_5Ca_ = 0.37, 95% CI = 0.36 – 0.39). Thus, both mutation render these receptors less sensitive to the blocking effects of Ca^2+^.

### Disease-associated variants display a broad and complex set of functional changes

The previous functional characterization of GluN2A^P552R^ and GluN2A^F652V^ variants suggested that both these variants are ‘gain-of-function’ mutations based on an observed increase in charge transfer and longer mean open durations ^5, 20^. However, given the large variability we observed for these mutants we undertook a systematic evaluation of the functional impact of the mutation relative to wild-type, by combining our gating and permeation data to estimate changes in responses to physiological pulses of glutamate.

We performed glutamate dose-response measurements for the two mutants. The response of GluN2A^F652V^ was unchanged relative to wild type: EC_50_, 4.3 ± 0.2 μM, vs 3.2 ± 0.1 μM, and *h,* 1.33 ± 0.07 vs. 1.31 ± 0.04). Therefore, for this mutant we used wild-type glutamate binding rates for our simulations. In contrast, and consistent with previous reports ^20^, GluN2A^P552R^ had substantially higher glutamate affinity (EC_50_, 0.51 ± 0.02 μM, *h* = 0.95 ± 0.03) (Figure 4a). Therefore, to simulate dynamic responses we needed to measure the glutamate binding rate for this variant. Given this receptor’s low P_o_, measuring the microscopic rates for glutamate association and dissociation would be intractable with a single-channel approach. Instead, we measured the apparent association and dissociation rate constants by fitting the rising and decay phases of the macroscopic current elicited with pulses of glutamate, sufficient to equilibrate the response (Figure 4b). We used the rates derived from fitting the model globally to all the single-channel records in each condition and the rates for glutamate association and dissociation, to simulated the macroscopic response to a prolonged exposure to saturating glutamate. We observed robust agreement between responses predicted by the models and the experimentally recorded whole-cell responses for wild-type, GluN2A^F652V^ and GluN2A^P552R^, indicated that the models capture the essential features of the reaction mechanism (Figure 4c; Table S2). Based on this result, we used the model to evaluate the potential impact of these variants on the response to several physiological-like stimuli.

**Figure 4.**
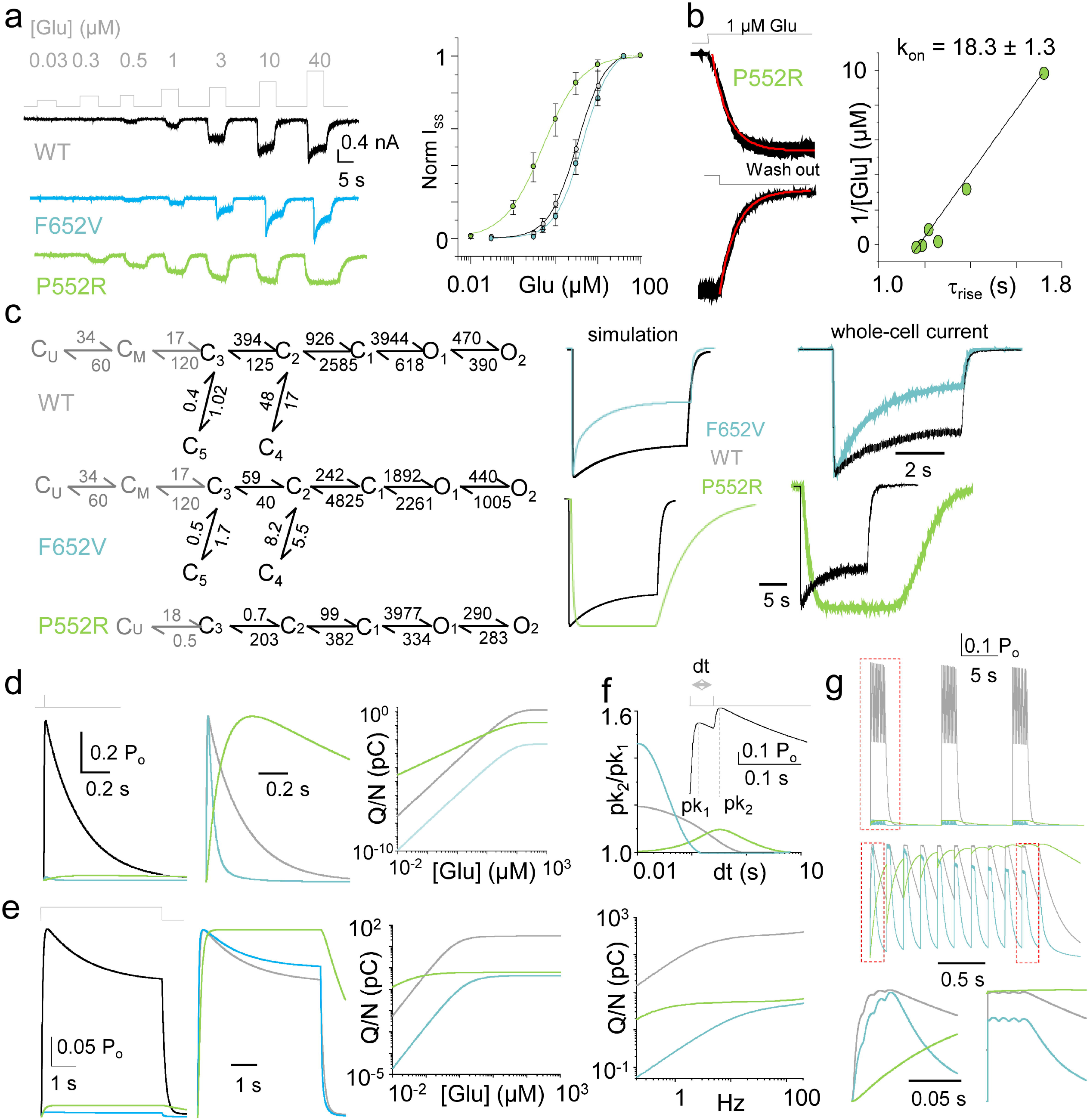
Disease-associated variants display complex functional changes. (**a**, *left*) Whole-cell currents evoked with several concentrations of glutamate in saturating glycine (0.1 mM); (*right*) Glutamate dose-dependence of the macroscopic steady-state current amplitude. (**b**, *left*) The rise and decay phases of the macroscopic current (black) recorded from the GluN1/GluN2A^P552R^ variant superimposed with fits to exponential functions (red). (*Right*) Glutamate dose-dependence of the rise time (circles) and fit to linear function (line) used to estimate the apparent glutamate binding rate. (**c**, *left*) Reaction mechanisms derived from fitting each model simultaneously to all single-channel recordings in each data set. (*Right*) Macroscopic current responses simulated with the respective kinetic models (*left*) and corresponding experimentally recorded whole-cell currents (*right*). (**d**) Simulated responses to a synaptic-like glutamate exposure (1 ms, 1 mM) predict drastic changes in peak current levels (left), time course (middle), and charge transfer. (**e**) Simulated responses to extrasynaptic-like glutamate exposure (5 s, 2 μM) predict complex changes in steady-state current levels (left), kinetics (middle), and charge transfer (right). (**f**) *Top*, Simulated responses to repetitive exposure to synaptic-like pulses predict complex changes in frequency-dependent facilitation. *Bottom*, Cumulative charge transfer over 60 sec of repetitive stimulation over varying physiologic frequencies. (**g**) *Top*, Simulation response to standard theta-burst stimulation. *Middle*, Expanded view of the normalized response to a single train of stimuli. *Bottom*, Expanded view of the normalized response to the first and last epoch of the first train of stimuli.

First, we simulated responses to a single synaptic-like pulse of glutamate (1 ms, 1 mM) in the continuous presence of glycine (Figure 4d). For a defined number of channels, both GluN1/GluN2A^P552R^ and GluN1/GluN2A^F652V^ produced substantially smaller currents than wild-type receptors. When normalized to the peak current amplitude, GluN2A^P552R^ current had slower rise and decay phases consistent with previous reports in recombinant and transfected neuronal systems ^20^. Further, when we quantified the total Ca^2+^-charge transferred per channel across the full simulation time and glutamate concentrations, we found that at all concentrations, GluN2A^F652V^ receptors consistently transferred less Ca^2+^ than wild-type receptors. In contrast, the Ca^2+^ transferred by GluN2A^P552R^ varied with glutamate concentration such that, at concentrations above 10 μM, it approached levels seen with wild-type receptors.

Extrasynaptic receptors likely experience chronic neurotransmitter exposure, initiate apoptotic pathways, and contribute to neurological disorders ^57^. Like with synaptic simulations, GluN2A^F652V^ consistently transferred 100-fold less charge than wild-type. In contrast, GluN2A^P552R^ transferred more charge at lower glutamate concentrations whereas at concentrations greater than 0.1 μM these receptors transferred less charge than wild-type. (Figure 4e).

We next evaluated the degree of potentiation in response to repetitive brief stimulation to mimic periods of high-frequency transmission. In wild-type receptors, the degree of potentiation dissipates as the duration between pulses lengthens ^41^. In the simulations performed here, relative to wild-type receptors, GluN2A^F652V^ receptors exhibited greater potentiation at shorter interpulse intervals. In contrast, GluN2A^P552R^ exhibited larger potentiation at longer interpulse intervals (Figure 4f). To assess channel sensitivity to a broad range of a physiologic range of stimuli frequency (0.2 – 200 Hz), we quantified the cumulative charge transfer per channel over 60 sec of stimulation. While both variants passed less total charge per minute compared to wild-type, the GluN2A^P552R^ variant exhibited less sensitivity over this range of stimuli frequencies.

Similarly, in response to a theta-like burst, within a given train of stimuli, the maximal GluN2AF652V current appeared within the first epoch whereas the maximal GluN2AP552R current appeared within later epochs of the protocol (Figure 4g). Thus, in addition to their primary deficit of reduced total charge transfer, different variants may exhibit delayed, time-dependent phenotypes depending on the physiological stimulation protocol.

## Discussion

Results reported here provide a systematic single-molecule characterization of the impact of two *Grin2A* disease variants on di-heteromeric receptor kinetics. We show that these two variants occur at sites critical to receptor gating whose interaction is necessary for normal function. Together with previous studies ^34, 58^, these results provide additional evidence for a role of direct interactions between residues on the GluN2A pre-M1 helix with those on the M3 helices of GluN2A and GluN1, as an intrinsic part of the gating machinery. Thus, the probability of channel opening and the stability of the open state are highly sensitive to atomic-level variations at these positions such that missense mutations will likely result in altered receptor responses. In addition, we found that structural variations at this interface also impact channel permeation properties.

Presently the full spectrum of biological functions of NMDA receptors is unknown. A major and critical role for NMDA receptors in the pathophysiology of the central nervous system is to produce a Ca^2+^-rich depolarizing current (excitatory) in the postsynaptic neuron. However, biological function depends critically on many other receptor capabilities, such as voltage-dependent Mg^2+^ block, glycine binding, etc. In addition, NMDA receptors are expressed at non-synaptic sites and in non-neuronal cells such as glia and are also present in tissues outside of the CNS, where their roles remain obscure.

### The multiple roles of the pre-M1 linker during gating

In all ionotropic glutamate receptors, the LBD is connected to the pore domain by three linkers D1-M1, D2-M3, and D2-M4. Identifying the precise motions of linkers during gating has been complicated by their high degree of freedom resulting in an inability to reliably resolve them structurally ^59–61^. Nevertheless, accumulating functional, genetic, and structural evidence in recent years has implicated linkers in mediating receptor function beyond serving as inert elements that tether domains. In NMDA receptors, loosening the D2-M3 linker with inserted glycine residues increases the opening latency after glutamate exposure demonstrating a role of mechanical tension in coupling agonist binding to the efficiency of pore opening ^62^. Linkers are also sites of drug binding ^63–65^. The GluN1 D2-M3 linker also provides a Ca^2+^ binding site necessary for enriching the NMDA receptor Ca^2+^ current ^66^. Both mutagenesis and swapping of linkers between receptor families have drastic effects on gating ^67, 68^. In congruence, our previous study revealed the coupling of the GluN2A D1-M1 linker with the GluN1 M3 helix can be perturbed by a single isomerization of a residue sidechain ^34^. Thus, the specific chemical properties of the linkers are as important for proper function as their length/mechanical properties.

In this study, we add to this pioneering literature by providing evidence for a direct coupling of the GluN2A D1-M1 linker with the GluN2A M3 helix which occurs late in the activation pathway, specifically at the C_2_-C_1_ and O_1_-O_2_ transitions (Figure 1). This interaction likely serves multiple roles including mediating efficient pore opening and stabilizing the open state and (Figure 5). We also note, that the GluN2A D1-M1 linker makes contacts with other structural elements including the GluN1 D2-M4 linker (Figure 1b), which may play a role in channel desensitization and provide insight into the profound effects the GluN2A^P552R^ variant exhibited on desensitization (Figure 2b). In tandem with previous findings, ^34, 58^, the D1-M1 linker makes several interactions and its rearrangement is a necessary and rate limiting step in the gating pathway. Decoupling the D1-M1 linker by inserting additional glycine residues is sufficient to perturb gating suggesting that agonist-induced tension in this linker is a rate limiting step in channel function. Furthermore, glycine insertion proximal to GluN2A P552 is sufficient to abolish receptor function ^58^. This supports both mechanical and chemical fidelity of the linker as necessary for function. This is because, in addition to loosening the mechanical tension of the linker, glycine insertion would also displace P552 relative to its native interacting partners, such as F652, thereby altering the efficiency of the chemical coupling of D1-M1 with M3

**Figure 5.**
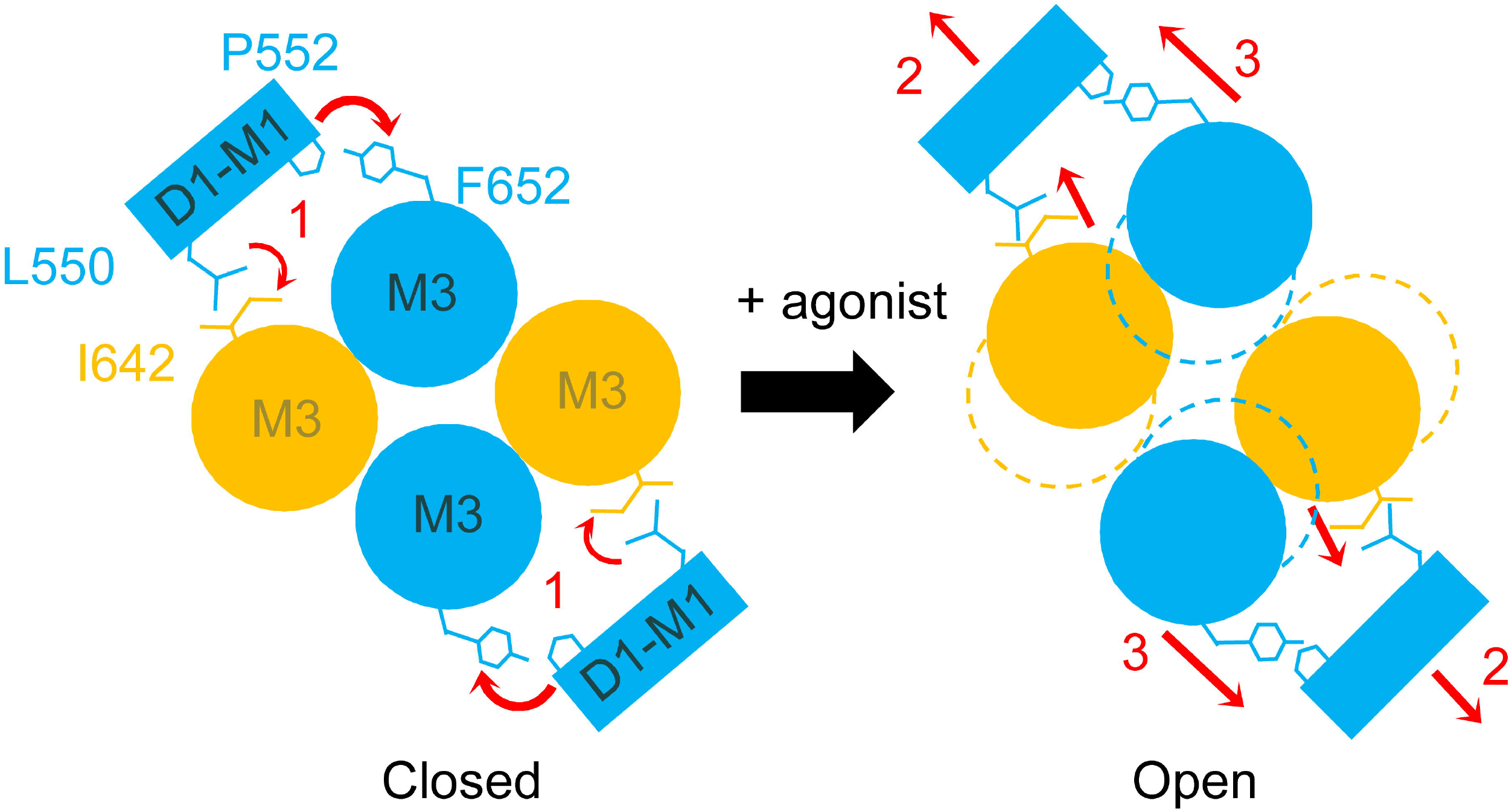
Proposed role of P552/F652 interaction in the gating reaction of NMDA receptors. (1) Residues on the D2-M1 linker of GluN2A subunits (blue), such as L550 and P552, interact directly with residues located on the gate-forming M3 helix of the same subunit (P552/F652) or of the adjacent GluN1 subunit (yellow) (L550/I642). (2) Agonist-triggered contraction of the LBD domain induces outward movement of the D2-M1 linker and causes (3) stabilization of the open M3 position.

Our results show strong coupling between D1-M1 linker and M3 at multiple steps in the activation pathway and thus suggest that the GluN2A D1-M1 linker is multifunctional. Consistent with this, its coupling with peripheral M4 helices ^58, 69^ occurs during gating and may underlie the fast component of gating. We have shown previously that the GluN2A D1-M1 linker can interact with the GluN1 M3 gate-forming helix to specifically stabilize its open position ^34^. Thus, specific motions of the D1-M1 linker may underlie different functionally distinct states in the gating reaction.

### Classification and treatment of NMDA receptor missense variants

In 2015, about 3.4 million people in the US were diagnosed with active epilepsy. Despite numerous pharmacologic treatment options available, only 44 % of those with active epilepsy report seizure control ^70^ consistent with reported rates of drug-resistant epilepsy ^71^. These numbers highlight the need for more defined pathophysiology and mechanism-targeted therapy. Epilepsy is a broad family of neurological disorders characterize by neuronal hyperexcitability. Numerous molecular mechanisms have been implicated in the pathogenesis of this heterogeneous family of disorders. Genetic studies have begun to shed light on this complexity by identifying *GRIN2A* mutations in severe forms of epilepsy ^2^, and *GRIN2A* variants associated with epileptic aphasias accounted for as much as 9 % of cases ^3^. Consistent with the causal association with disease pathogenesis, variant distribution across GluN2A domains was correlated with clinical/electrophysiological phenotype ^72^. Within this study, however, phenotypic variations existed among individuals with the same variant and it was recognized that the complex functional alterations caused by a *GRIN2A* variant cannot be reduced to a binary description such as loss- or gain-of-function ^72^.

The functional impact of a single missense variant may have unique manifestations in different physiological conditions. For a multifunctional protein with numerous physiological outputs such as the NMDA receptor, a single mutation may exhibit both gain- and loss-of-function properties under different stimuli for different physiological outputs (Figure 4). This may explain recent observations showing that disease variants on GluN2A and GluN2B classified as either gain- or loss-of-function based on microscopic parameters behavior largely indistinguishably *in vivo* ^32, 33^. This will impact how variants should be classified and, thus, how to tailor treatment to individuals harboring a specific mutation. A previous report have classified GluN2A^F652V^ variant as gain-of-function based on a single gating parameter ^5^. However, we observe that a single variant can exhibit either gain- or loss-of-function depending on the parameter measured (Table S3). Here, we note a correlation between the extent of contacts between P552 and F652 and channel open duration, which supports a key role for this interaction in defining the stability of the open channel conformations. The GluN2A^F652V^ variant has a reduced contact surface area and exhibits a shorter mean open duration, whereas GluN2A^P552R^ increases the extent of contacts and exhibits longer mean open durations (Figure 2). Further N2A^P552A^, which only moderately reduces the number of contacts, has a less substantial impact on gating (Figure 1)^73^. Therefore, when interpreting the functional effect of a variant, it is important to consider the physicochemical properties of the variant not only the site of the missense substitution.

This has important implications when designing treatment regimens for patients with specific mutations. Furthermore, different disease-variants, even at distant sites of action, can have profound influence on affinity and mechanism of drug action. GluN2A^P552R^ receptors are among the variants reported to have substantially reduced affinity and distinct mechanism of action of an Alzheimer’s drug derivative ^74^. Therefore, not only is a comprehensive functional and kinetic analysis of a disease variant is necessary prior to designing and testing therapeutics, but also an investigation of the effect of a specific variant on the pharmacodynamics of existing drugs.

### The case for design of state-specific pharmacological modulation

Given the causal association of NMDA receptor variants with epilepsy pathogenesis, pharmacological targeting of NMDA receptors holds promising therapeutic potential. However, because NMDA receptors are indispensable to synaptic physiology, global modulation of receptor function can result in neurotoxicity. Therefore, there is interest in designing modulators that target specific receptor functions ^75^. This may in part underlie the success of memantine in the clinical treatment of neurological disorders. For example, recent evidence suggests that memantine can specifically modulate Ca^2+^-dependent inhibition of channels ^76^. This strategy requires knowledge of the precise structural elements and their dynamics which underlie specific receptor functions. This remains a large knowledge gap in the field. Our approach of using structural model-guided mutagenesis with single-molecule derived kinetic characterization begins to bridge this gap.

Several recent studies suggest that modulators targeted to linker regions can independently control gating and permeation ^65^. This is consistent with our observation that mutations in this region exhibited lower unitary amplitudes and altered Ca^2+^ permeation ^34^ (Figure 3). In addition, subtle changes in the chemical structure of modulators acting in this region can change a negative allosteric modulator to a positive allosteric modulator ^64^. This is consistent with our observation that a single isomerization mutation that alters the interaction between the GluN2A D1-M1 linker and the GluN1 M3 helix is sufficient to decrease function ^34^. The use of empirical kinetic models to map precise structural elements to specific receptor functions provides a workflow for the design of function-specific pharmacological modulators. The use of kinetic mechanism-based pharmacological targeting represents a new avenue for precision medicine ^77^.

Although the number of identified variants continues to rise, only few have been characterized functionally and remains unknown which reported functional changes cause pathology. Given that several functional attributes of NMDA receptors are critical for the normal physiologic response and thus for their biological role, it remains unknown how any variant affects the pathophysiology of the cells in which it is expressed and further the behavior of the individual patient. Most neuropsychiatric conditions that are currently associated with NMDA receptor dysfunction are complex and of unresolved etiology such as: epilepsy, language disorders, motor disorders, learning disorders, autism, attention deficit hyperactivity disorder, developmental delay, and schizophrenia. Therefore, the field will require more in depth understanding of receptor operation before rendering rational therapeutic strategies.

## Acknowledgements

We thank Jamie Abbott, PhD for helpful critiques and technical support in cell culture and molecular biology. This study was funded by NIH R01NS108750 and R01NS097016 to GKP. GJI and GKP conceived the study. HW and WZ performed and analyzed MD simulations. GJI, BL, and BS performed and analyzed electrophysiology experiments. GJI, BL, and GKP prepared figures and wrote the manuscript.

## Conflict of Interest

All authors declare no financial or non-financial conflicts of interest with the content of this article.

Supplementary information is available at MP’s website.

## Supplementary Figures

**Figure S1.**
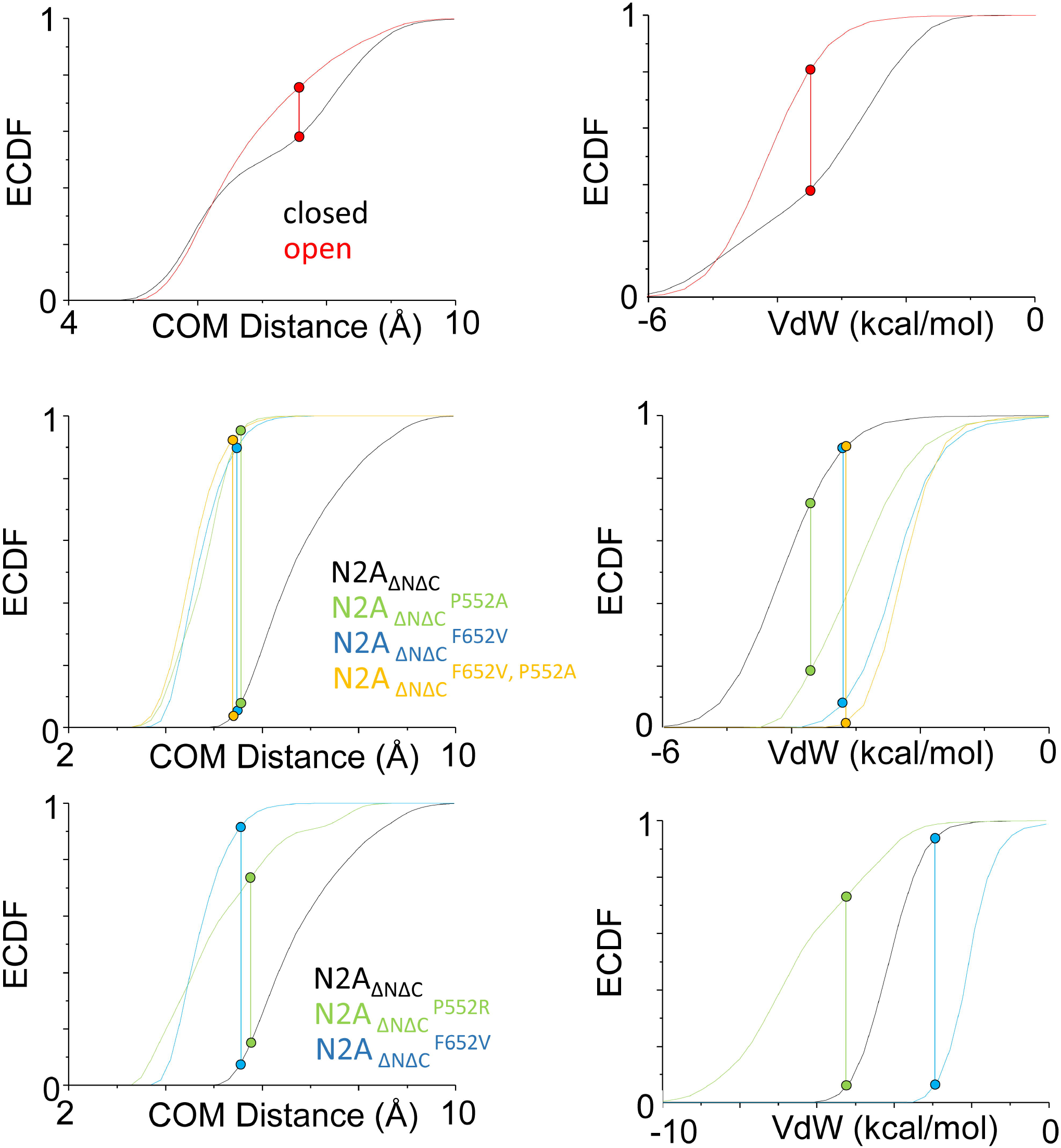
Empiric cumulative distribution functions (ECDF) of residue sidechain center-of-mass (COM) distance and van der Waals (VdW) contact energy during MD simulation of the active/open structure for each construct. Data points and dashed lines indicate Kolmogorov-Smirnov test statistic for each construct.

## Supplementary Tables

**Table S1:** Summary of single-channel gating parameters

|  | n | i (pA) | P<sub>o</sub> | MCT (ms) | MOT (ms) | Record time (min) | Total events |
| --- | --- | --- | --- | --- | --- | --- | --- |
| GluN2A | 11 | 8.9 ± 0.4 | 0.46 ± 0.05 | 6 ± 1 | 3.9 ± 0.4 | 360 | 2,608,269 |
| GluN2A<sup>P552R</sup> | 16 | 4.4 ± 0.1* | 0.02 ± 0.01* | 479 ± 98* | 5.5 ± 0.8 | 293 | 56,838 |
| GluN2A<sup>F652V</sup> | 14 | 5.6 ± 0.2* | 0.01 ± 0.00* | 92 ± 18* | 0.7 ± 0.1* | 355 | 307,378 |

\* $p < 0.05$ relative to WT (Student's $t$ test)

**Table S2:** Summary of MIL exponential fits

|  | Closed Components |  |  |  |  | Open Components |  |
| --- | --- | --- | --- | --- | --- | --- | --- |
| | $\tau_3$ (A<sub>3</sub>) | $\tau_2$ (A<sub>2</sub>) | $\tau_1$ (A<sub>1</sub>) | $\tau_4$ (A<sub>4</sub>) | $\tau_5$ (A<sub>5</sub>) | $\tau_1$ (A<sub>1</sub>) | $\tau_2$ (A<sub>2</sub>) |
| GluN2A | 0.16 ± 0.01<br>(0.35 ± 0.04) | 1.24 ± 0.07<br>(0.29 ± 0.04) | 4.4 ± 0.4<br>(0.34 ± 0.04) | 24 ± 5<br>(0.015 ± 0.003) | 2947 ± 429<br>(0.002 ± 0.000) | 0.12 ± 0.01<br>(0.11 ± 0.02) | 3.0 ± 0.5<br>(0.47 ± 0.04) |
| GluN2A<sup>P552R</sup> | 0.26 ± 0.02*<br>(0.79 ± 0.05*) | 3.5 ± 0.7*<br>(0.13 ± 0.06) | 167 ± 89<br>(0.016 ± 0.003*) | 9514 ± 1881*<br>(0.10 ± 0.01*) | -- | 1.5 ± 0.4*<br>(0.22 ± 0.04*) | 9.4 ± 1.3*<br>(0.69 ± 0.04*) |
| GluN2A<sup>F652V</sup> | 0.15 ± 0.01<br>(0.14 ± 0.01*) | 8.4 ± 1.2*<br>(0.26 ± 0.03*) | 39 ± 5*<br>(0.49 ± 0.03*) | 384 ± 107*<br>(0.08 ± 0.02*) | 2393 ± 590<br>(0.029 ± 0.004*) | 0.37 ± 0.03*<br>(0.63 ± 0.06*) | 1.20 ± 0.09*<br>(0.35 ± 0.05*) |

\* $p < 0.05$ relative to WT (Student's $t$ test)

**Table S3:** Summary of phenotype classification by parameter*

|  | GluN2A<sup>P552R</sup> | GluN2A<sup>F652V</sup> |
| --- | --- | --- |
| P<sub>o</sub> | LOF | LOF |
| P<sub>o,burst</sub> | GOF | LOF |
| MOT | - | LOF |
| MCT | LOF | LOF |
| $\gamma_{Na}$ | LOF | LOF |
| $\gamma_{2Ca}$ | - | - |
| $\gamma_{2Ca}/\gamma_{Na}$ | GOF | GOF |
| $\gamma_{5Ca}/\gamma_{Na}$ | GOF | GOF |
| P<sub>5Ca</sub>/P<sub>Na</sub> | LOF | - |
| P<sub>10Ca</sub>/P<sub>Na</sub> | LOF | - |
| Glu EC<sub>50</sub> | GOF | - |
| $\tau_D$ | GOF | LOF |

\* classification based on predicted relative effect on Ca<sup>2+</sup> flux

## Notes

### Competing Interest Statement

The authors have declared no competing interest.

